# GEMC1 and MCIDAS interactions with SWI/SNF complexes regulate the multiciliated cell-specific transcriptional program

**DOI:** 10.1101/2022.12.02.518887

**Authors:** Michael Lewis, Berta Terre, Philip A. Knobel, Tao Cheng, Hao Lu, Camille Stephan-Otto Attolini, Jordann Smak, Etienne Coyaud, Isabel Garcia-Cao, Jessica Querol, Gabriel Gil-Gomez, Gabrielle Piergiovanni, Vincenzo Costanzo, Sandra Peiró, Brian Raught, Haotian Zhao, Xavier Salvatella, Sudipto Roy, Moe Mahjoub, Travis H. Stracker

## Abstract

Multiciliated cells (MCCs) project dozens to hundreds of motile cilia from their apical surface to promote the movement of fluids or gametes in the mammalian brain, airway or reproductive organs. Differentiation of MCCs requires the sequential action of the Geminin family transcriptional activators, GEMC1 and MCIDAS, that both interact with E2F4/5-DP1. How these factors activate transcription and the extent to which they play redundant functions remains poorly understood. Here, we demonstrate that the transcriptional targets and proximal proteomes of GEMC1 and MCIDAS are highly similar. However, we identified distinct interactions with SWI/SNF subcomplexes; GEMC1 interacts primarily with the ARID1A containing BAF complex while MCIDAS interacts primarily with BRD9 containing ncBAF complexes. Treatment with a BRD9 inhibitor impaired MCIDAS-mediated activation of several target genes and compromised the MCC differentiation program in multiple cell based models. Our data suggest that the differential engagement of distinct SWI/SNF subcomplexes by GEMC1 and MCIDAS is required for MCC-specific transcriptional regulation and mediated by their distinct C-terminal domains.

## Introduction

In most cell types, DNA replication is limited to once per cell cycle to avoid endoreduplication that can trigger polyploidy, replication stress and cell death. Several mechanisms restrain re-replication, including the phosphorylation or proteolytic regulation of pre-replication complex (Pre-RC) components, the activity of checkpoint kinases and the inhibition of CDT1-dependent origin licensing by physical interactions with Geminin^1^. Geminin contains a central coiled-coil (CC) domain that is required for its dimerization and interaction with CDT1. Similar CC domains were identified in the Geminin coiled-coil containing protein 1 (GEMC1, encoded by *GMNC*) and MCIDAS/Multicilin (encoded by *MCIDAS*) and shown to be critical for their functions^2–6^. Both GEMC1 and MCIDAS have been implicated in the regulation of DNA replication through antagonistic interactions with Geminin^2,3,7–9^.

GEMC1 and MCIDAS also play critical roles in the transcriptional differentiation of multiciliated cells (MCCs), specialized epithelial cells that project dozens to hundreds of motile cilia from their apical surfaces and play crucial roles in the olfactory/respiratory, reproductive, renal and central nervous systems of many vertebrates^5,9–13^. To activate transcription of this unique pathway, both proteins interact with E2F4/5-DP1 through a conserved C-terminal domain absent in Geminin, dubbed the TIRT domain due to a repeating amino acid motif in MCIDAS^5,9,12^. This domain is required for the transcriptional activity of both GEMC1 and MCIDAS, and mutations in the TIRT domain of MCIDAS have been implicated in the rare ciliopathy Reduced Generation of Multiple Motile Cilia (RGMC) that is characterized by the loss or dysfunction of motile cilia on MCCs, causing hydrocephaly, defects in respiratory physiology and infertility^14,15^. The TIRT domains of GEMC1 and MCIDAS convey differential affinity for E2F proteins, with GEMC1 binding preferentially to E2F5 and MCIDAS to both E2F4 and E2F5^13^.

The expression of *Gemc1* precedes that of *Mcidas* in the developing mouse brain, suggesting that GEMC1 acts upstream of MCIDAS^11^. Supporting this, mice lacking GEMC1 have no identifiable MCCs in any tissues, while mice lacking MCIDAS have columnar MCC-like cells that are positive for MCC markers, including FOXJ1 and TP73, but fail to amplify centrioles and generate motile cilia^5,12,13,16^. GEMC1 activates several genes critical for MCC generation, including *MCIDAS*, *CDC20B* and *CCNO* that reside in the same chromosomal region, as well as key transcription factors such as MYB, FOXJ1 and TP73^5,6,12,16,17^.

Both GEMC1 and MCIDAS share numerous protein-protein interactions and play similar roles in MCC differentiation, but their level of redundancy remains unclear. To understand their relative molecular functions, we compared the transcriptional targets and proximal interactomes of GEMC1 and MCIDAS. Our results revealed that both proteins activate a core set of overlapping genes, but also have distinct transcriptional targets. The proximal interactomes of each protein were strikingly similar and composed almost exclusively of proteins related to transcription, including almost all transcription factors implicated in multiciliogenesis to date^18^. Notable among these is the SWI/SNF (switching/sucrose non-fermenting) chromatin remodeling complex that functions with E2F4/5 in transcriptional regulation^19^. We demonstrate that TIRT domains of GEMC1 and MCIDAS confer preferential interactions with distinct SWI/SNF sub-complexes and the depletion or inhibition of SWI/SNF components impairs their ability to activate transcription. Finally, inhibition of BRD9, a specific subunit of the ncBAF SWI/SNF complex impaired multiciliogenesis in mouse tracheal epithelial cells (mTECs) cultured at air liquid interface (ALI), as well as glioma cells overexpressing MCIDAS. Together, our results provide an overview of the core transcriptional machinery associated with GEMC1 and MCIDAS and demonstrate a critical role for SWI/SNF in GEMC1/MCIDAS-mediated transcriptional regulation during multiciliogenesis.

## Materials and Methods

### Cell culture and transfection

AD-293 (Agilent), HEK-293T, cells and Hela cells were cultured in Dulbecco’s Modified Eagle Medium (Gibco) with 10% FBS (Hyclone). DBTRG-05MG cells (DSMZ) were cultured in RPMI-1640 with 10% FBS. Cells were routinely tested for mycoplasma and found negative (ATCC). For transient transfections, cells were seeded in either 10 cm or 6 well plates at 70% confluence and transfected with 10 µg or 3 µg of plasmid respectively, with polyethylenimine (Polysciences) for AD-293 or Lipofectamine 2000 (ThermoFisher Scientific) for Hela cells. The medium was changed 8 hrs post-transfection and cells were collected after 48 hrs.

### Reagents and antibodies

All chemicals and reagents were purchased either from Sigma-Aldrich (Merck) or Thermo Fisher Scientific (Invitrogen). Commercial antibodies used are listed in Supplementary Table S1.

### RNA purification and microarrays

AD-293 cells were transfected as described in 10 cm plates, cells were washed in PBS and lysed in Trizol (Invitrogen) 48 hours post-transfection. RNA was purified using the PureLink RNA mini kit (Thermo-Fisher). Microarrays were performed by the IRB Functional Genomics Facility. Briefly, cDNA was generated from 25 ng of RNA using the TransPlex Complete Whole Transcriptome Amplification Kit (Sigma) and 16 cycles of amplification. 8 µg of cDNA was fragmented and labeled using GeneChip Human Mapping 250K Nsp Assay Kit (Affymetrix), according to manufacturer’s instructions. cDNA was hybridized to the Genechip Primeview Human Array (Affymetrix) for 16 hours at 45°C in a GeneChip Hybridization oven 645 (Affymetrix). Washing and staining of microarrays was performed using a GeneChip Fluidics Station 450 (Affymetrix) and arrays were scanned with GeneChip scanner GSC3000 (Affymetrix). Affymetrix GeneChip Command Console software (AGCC) was used to acquire GeneChip images and generate CEL files for analysis. Arrays were imported to R applying RMA background correction and probe summarization. For each tissue, fold changes between conditions were computed after mean and variance normalization and centering using a generalized additive model (GAM). An empirical Bayesian partial density model was used to identify differentially expressed genes. Gene set enrichment analysis (GSEA) was performed using all genes in the array sorted by fold change computed in each tissue using GOSlim datasets^20^.

### Western blotting assays

For western blotting, cells were collected and lysed in RIPA buffer (1% NP-40, 0.1% sodium dodecyl sulfate (SDS), 50 mM Tris–HCl pH 7.4, 150 mM NaCl, 0.5% sodium deoxycholate, 1X proteinase inhibitor cocktail (Roche)) on ice. Samples were sonicated using a Bioruptor XL sonicator (Diagenode) for 15 min with 15 sec intervals and centrifuged at 4°C for 15 min at 1300 rpm. The proteins were quantified and resolved on 8% SDS-polyacrylamide gels with 1% (v/v) β-mercaptoethanol, 0.01% (w/v) Bromophenol blue and then transferred onto nitrocellulose membrane (Bio-Rad) at 100V for 1 hour. The membrane was blocked with 5% milk powder (Bio-Rad) then incubated with specific antibodies at 4°C overnight. Following incubation with secondary antibodies, immunoblots were visualized LI-COR Odyssey CLx system (LI-COR Biotechnologies). All uncropped blots are provided in the Supplementary Material.

### BioID analysis of proximity interactions

Human GEMC1 or MCIDAS were amplified from previously described expression vectors^5^ using forward primers containing *Asc*I and reverse primers containing *Not*I restriction sites (GEMC1-Asc1-F-AAGGCGCGCCATGAACACCATTCTGCCTT; GEMC1-NotI-R- TTGCGGCCGCCTAACTGGGGACCCAGCGGAACT; MCIDAS-Asc1-F- AAGGCGCGCCATGCAGGCGTGCGGGGGCGGC; MCIDAS-NotI-R- TTGCGGCCGCCTAACTGGGGACCCAGCGGAACT) using KOD Hot Start DNA Polymerase (Millipore) and cycling conditions recommended from the manufacturer (polymerase activation at 95 °C for 2 min, denaturation at 95 °C for 20 s, annealing at 55 °C for 10 s and extension at 70 °C for 50 s, repeated for 40 cycles). PCR products were purified using the PureLink Quick Gel Extraction Kit (Invitrogen) and cloned into pCR2.1-TOPO vector (Invitrogen). Top10 competent *E. coli* cells (Invitrogen) were transformed with pCR2.1-GEMC1/MCIDAS and colonies were selected with carbenicillin. Constructs were verified by restriction digestion and sequencing (Macrogen) with primers for the TOPO vector (T7 Promoter-F and M13-R). Afterwards, GEMC1 or MCIDAS were cut from the pCR2.1-TOPO vector by restriction digest with *Asc*I (New England BioLabs (NEB), Ipswich, MA, USA) and *Not*I-HF (NEB), purified using the PureLink Quick Gel Extraction Kit (Invitrogen) and ligated into pcDNA5/FRT/TO-N-FLAG-BirA* using Quick Ligation Kit (NEB). Top10 competent *E. coli* cells (Invitrogen) were transformed with pcDNA5/FRT/TO-N-Flag-hBirA*-GEMC1/MCIDAS vector and carbenicillin selected. The constructs were confirmed by restriction digestion with *Asc*I (NEB) and *Not*I-HF (NEB) and sequencing (Macrogen).

AD-293 cells were seeded and transfected the next day with either pcDNA5/FRT/TO-N-FLAG-hBirA*-GEMC1/MCIDAS or pcDNA5/FRT/TO-N-FLAG-hBirA* using PEI (Sigma-Aldrich) ±50 *μ*M biotin (IBA GmbH; 2-1016-002). For mass spectrometry, 5X15 cm^2^ plates per condition were harvested 24 h after transfection by scraping cells into PBS, washing two times in PBS and snap freezing on dry ice. Cell pellets were lysed in 5 ml modified RIPA buffer (1% TX-100, 50 mM Tris-HCl, pH 7.5, 150 mM NaCl, 1 mM EDTA, 1 mM EGTA, 0.1% SDS, 0.5% sodium deoxycholate and protease inhibitors) on ice, treated with 250 U benzonase (Millipore) and biotinylated proteins were isolated using streptavidin-sepharose beads (GE Healthcare). Proteins were washed in ammonium bicarbonate and digested with trypsin. Mass spectrometry and data analysis using SAINTexpress was performed as described previously^21,22^ and dot plot figures generated using ProHits-viz (https://prohits-viz.org).

Small scale BioID-AP followed by western blotting was carried out as described above, but using 3X15 cm^2^ plates per condition and, following affinity purification with streptavidin-sepharose beads (GE Healthcare), samples were centrifuged briefly and the supernatant aspirated. Samples were resuspended in 100ul-150ul RIPA buffer and boiled at 95°C for 10 minutes. Lysates were separated on SDS-PAGE gels and transferred for western detection as described previously.

### Geminin family Hybrid cloning

Hybrid constructs (domains detailed in Figure 3) were synthesized by Genescript into pcDNA3.1(+)-N-DYK. All hybrid constructs were tested for expression and cloned into the pcDNA5/FRT/TO-N-FLAG-hBirA* using restriction enzymes: 5’ *Kpn*I (NEB) and 3’ *Not*I (NEB) before being tested by sequencing (Macrogen) to confirm ligation.

**Figure 1:**
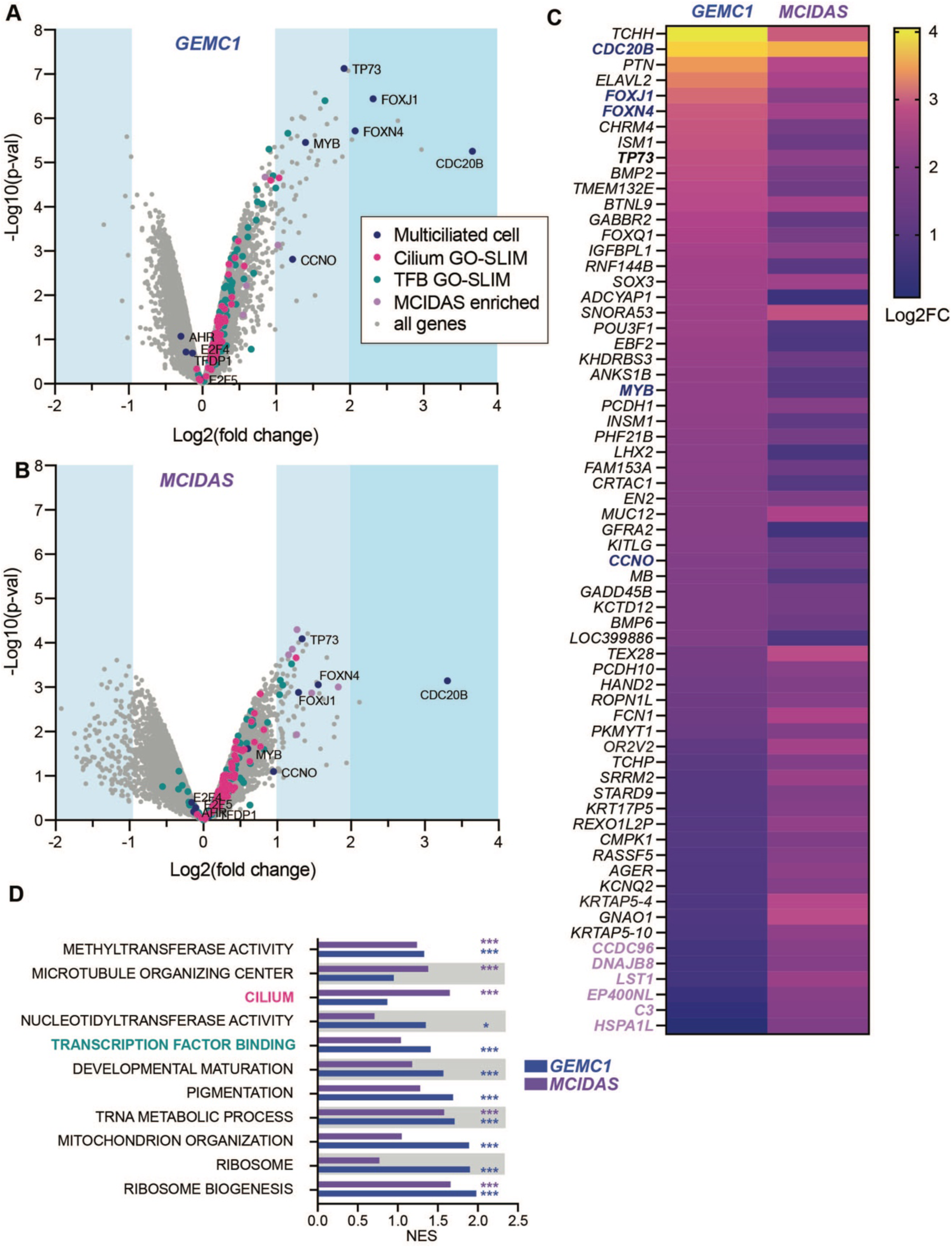
GEMC1 and MCIDAS overexpression activates overlapping and distinct target genes. **A.** Volcano plots (-Log10 p-value vs Log2 fold change) of microarray analysis of gene expression following transient transfection of *GEMC1* and **B.** *MCIDAS* in AD-293 cells. Genes in the Cilium and Transcription Factor Binding GO-SLIM categories (TFB = Transcription factor binding), genes implicated specifically in multiciliated cells or genes enriched with *MCIDAS* over *GEMC1* are indicated by color (legend in A, applies to A-D). Full details in Supplementary Tables S2 and S3. **C.** Heatmap comparing expression (Log2 fold change=Log2FC) of most upregulated genes following either *GEMC1* or *MCIDAS* expression. Color coding of gene categories is applied as in A. **D.** Gene set enrichment analysis of the gene expression data using GO-SLIM categories. The nominal enrichment score (NES) for each category with either GEMC1 or MCIDAS is shown. Details in Supplementary Table S3.

**Figure 2:**
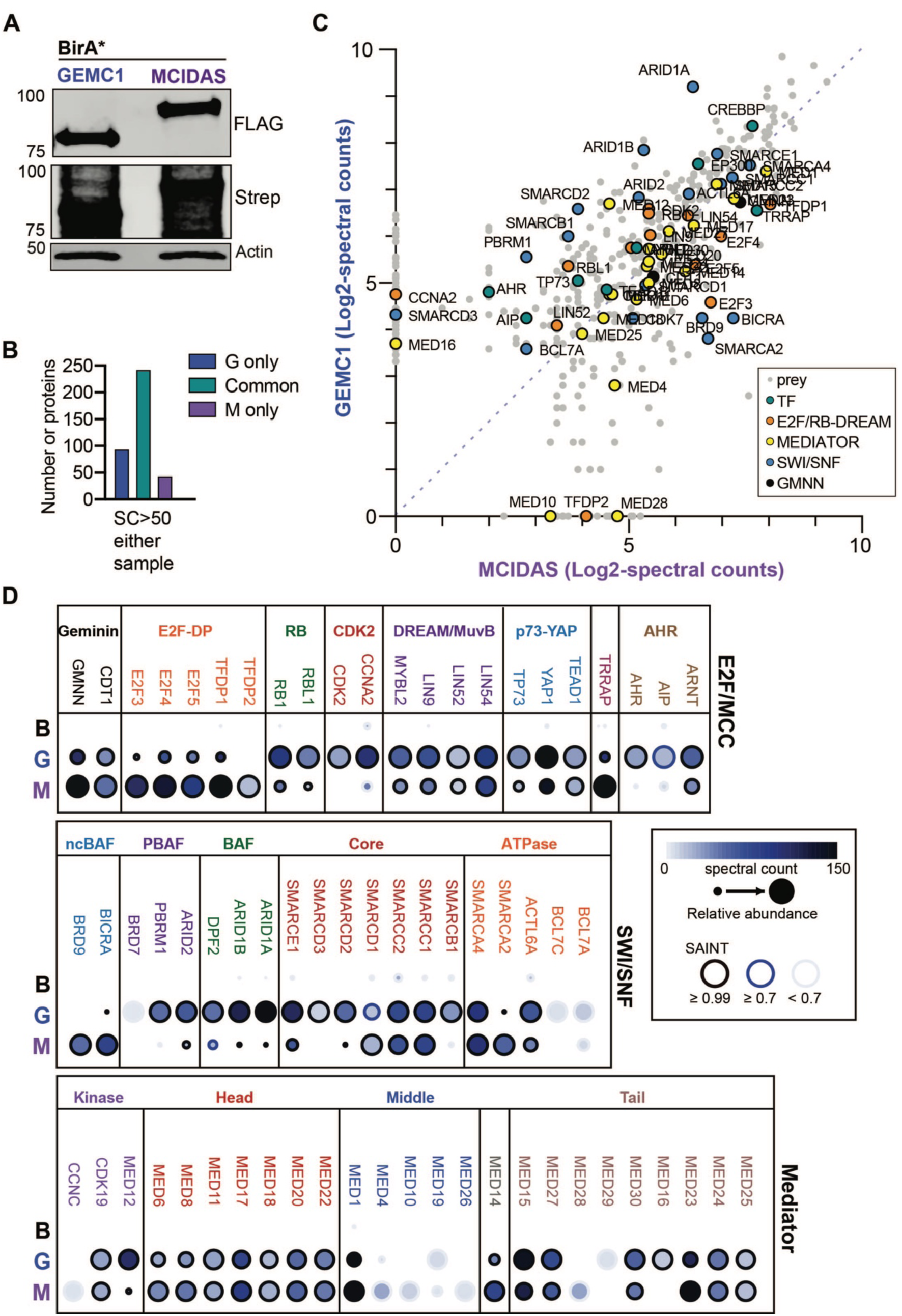
The proximal interactomes of GEMC1 and MCIDAS. **A.** Western blots from lysates of AD-293 cells transfected with FLAG-BirA*, FLAG-BirA*-GEMC1 or FLAG-BirA*-MCIDAS. Expression of baits in cells with or without biotin supplementation is shown using the FLAG antibody (bottom panel) and labeling in biotin supplemented samples with Strep-HRP (top panel). **B.** Graph of protein number unique or common to each sample using a cutoff >50 spectral counts (SC) with a SAINT score of >0.7 in either one of the samples. **C.** Scatter plot of peptides identified in GEMC1 or MCIDAS samples (Log2-spectral counts of 2 technical replicates). All peptides with a SAINT score of >0.7 in one of the samples are shown (Full list in Supplementary Table S4). Specific subsets are highlighted including key transcription factors (TF), components of the E2F or DREAM complexes, SWI/SNF and Mediator related proteins. **D.** Dot plot depicting relative abundance and spectral counts for selected proteins. B=BirA*, G=GEMC1 and M=MCIDAS. Proteins are grouped by those involved in E2F or MCC differentiation (top row), components of SW/SNF complexes (middle row) and components of the Mediator complex (bottom row).

**Figure 3:**
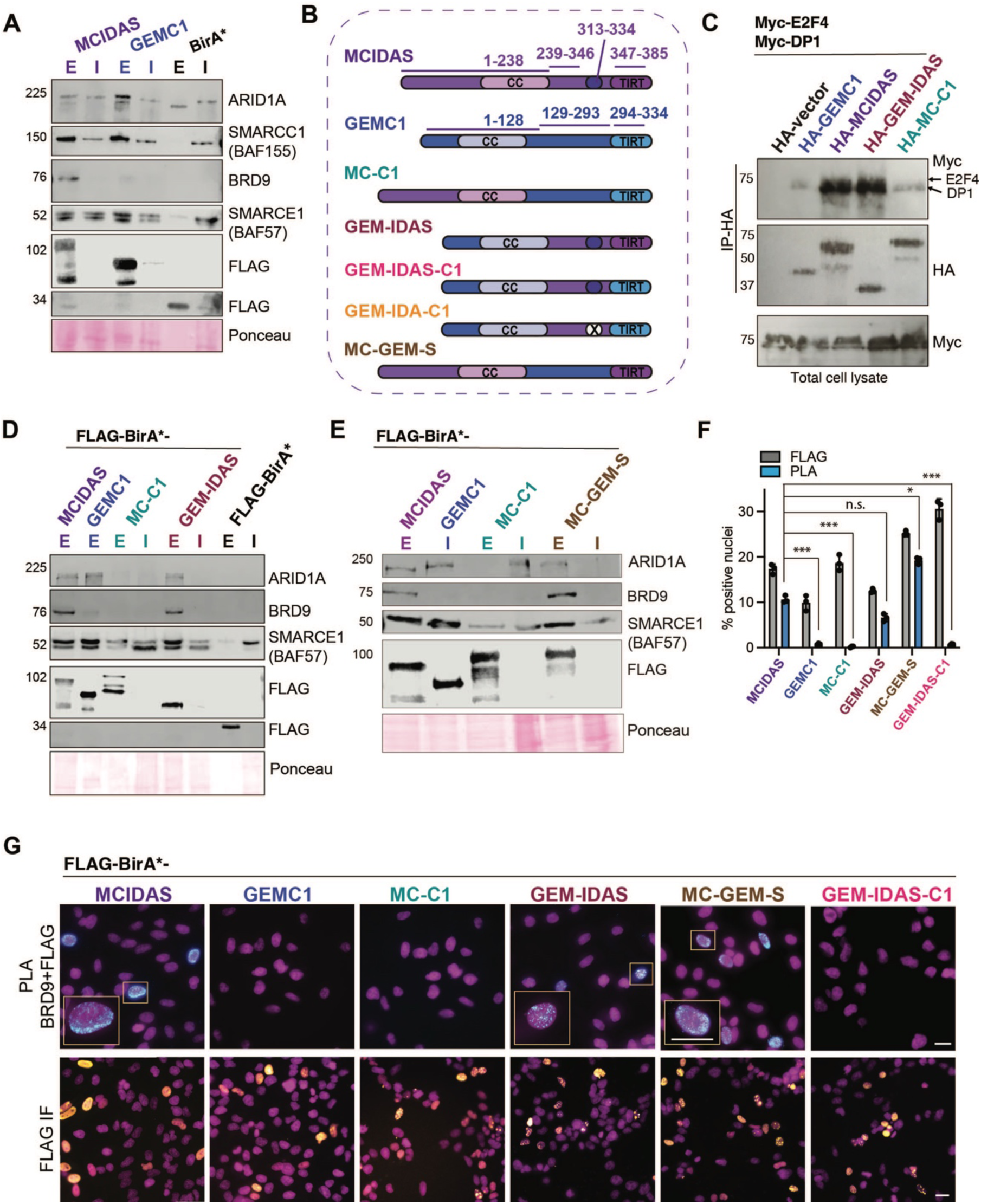
Specific interactions with SWI/SNF subcomplexes. **A.** BioID-AP western blots for ARID1A, BRD9 and core SWI/SNF subunits SMARCE1 (BAF155) and SMARCC1 (BAF57) demonstrate the relative specificity for BAF and ncBAF complexes. All uncropped blots are shown in Supplementary Figure S1. **B.** Schematic illustration of the structure of GEMC1-MCIDAS hybrid proteins. **C.** Co-immunoprecipitation experiments demonstrate enhanced E2F4-DP1 interactions with the MCIDAS C-terminus. HEK293T cells were transfected with HA-tagged vector, GEMC1, MCIDAS, GEM-IDAS and MC-C1, Myc-tagged DP1 and Myc-E2F4. Lysates were immunoprecipitated with anti-HA antibodies and westerns carried out for Myc and HA following transfer to PVDF. Total lysates are shown blotted for Myc. Data presented is representative of 2 biological replicates. **D, E.** BioID-AP westerns blots for ARID1A, BRD9 and core SWI/SNF components following expression of hybrid proteins (see B for schematic). Ponceau staining shown for loading and transfer control. **F.** Quantification of the percentage of transfected HeLa cells and PLA positive cells is shown in right panel. Mean (bar) with standard deviation and values of individual experiments (circles) are shown. For statistical analysis, an unpaired t-test on the PLA+ cells normalized to transfection efficiency was performed from 3 independent experiments. MCIDAS vs GEMC1, p=<1e-06(***), MCIDAS vs MCI-C1, p=<1e-06(***), MCIDAS vs GEM-IDAS, p=.836(n.s), MCIDAS vs MC-GEM-S, p=.0074(*), MCIDAS vs GEM-IDAS-C1, p=<1e-06(***). **G.** Representative immunofluorescence (IF) images of HeLa cells transfected with the indicated FLAG-BirA* tagged proteins (bottom panel, FLAG in yellow and DAPI (DNA) in magenta) and PLA-IF (top panels, PLA in cyan and DAPI (DNA) in magenta). Scale bar = 20 μM.

### Co-immunoprecipitation

Co-immunoprecipitation assays followed by western blot were performed as previously described. Briefly, GEM-IDAS and MC-C1 hybrid cDNAs were PCR amplified using forward primer containing *Hind*III and reverse primer containing *Not*I restriction enzyme sites (GM-F-HindIII: GATCGATCAAGCTTACCATGAACACCATTCTGCC; GM-R-NotI: GATCGATCGCGGCCGCCTACTGGGGACCCAGCGGAA; MG-F-HindIII: GATCGATCAAGCTTACCATGCAGGCGTGCGGGGGC; MG-R-NotI: GATCGATCGCGGCCGCCTAGACTGCTTAGGGACCCA) and cloned into the pXJ40 vector with one HA-tag at the N-terminus. Combinations of plasmids were co-transfected into HEK293T cells (3 µg plasmid, per 10 cm dish) using Lipofectamine 2000 (Thermo Fisher Scientific). 24 hrs post-incubation, cells were lysed in 800 µl of RIPA buffer (Thermo Fisher Scientific) supplemented with EDTA-free Protease Inhibitor Cocktail (Roche, 11836170001). Cell lysates were sonicated briefly, spun down and an aliquot taken for total cell lysate control. Remaining lysate was rotated O/N with 25 µl of Protein-A-agarose beads (Roche) and 2 µg of mouse anti-HA antibody (Supplementary Table S1). Beads were washed four times in RIPA buffer and boiled in 50µl of 1× SDS loading buffer. Total cell lysates (15µl, 1%) and IP (15µl, 30%) were resolved with SDS-PAGE gels, transferred to PVDF membranes, blocked in 2% BSA and probed with relevant primary antibodies for western detection.

### Quantitative real time-PCR (RT-qPCR)

Transfected AD-293 cells were collected after two cold PBS washes by scraping into Tri-Reagent. RNA was isolated by chloroform extraction followed by centrifugation, isopropanol precipitation, 2 washes in 75% ethanol and resuspension in DEPC-treated water. Nucleic acid quantification was performed with a Nanodrop 8000 Instrument (ThermoFisher Scientific). cDNA was generated using 0.5-1μg of total RNA and a High Capacity RNA-to-cDNA Kit (Applied Biosystems). RT-qPCR was performed using the comparative CT method and a StepOne Real-Time PCR System (Applied Biosystems). Amplification was performed using Power SYBR Green PCR Master Mix (Applied Biosystems) or TaqMan Universal PCR Master Mix (Applied Biosystems). All assays were performed in duplicate. For TaqMan assays, *mActB* probe was used as an endogenous control for normalization and a specific Taqman probe was used for mouse or human *TP73* (Hs01056231_m1), *CCNO* (Hs004389588_91), *FOXJ1* (Hs00230964_m1), and *TP73* (Hs01056231_m1). Primers used for SYBR Green assays (Sigma-Aldrich) can be found in Table S4.

### Proximity Ligation Assay (PLA)

Protein-protein interactions were studied using a Duolink In Situ Orange Starter Kit Mouse/Rabbit (Sigma DUO92102) following the manufacturer’s protocol. Briefly, transfected Hela cells were seeded in coverslips and cultured overnight. Slides were washed with cold 1XPBS and fixed in 4% paraformaldehyde for 15 min, permeabilized using 0.1% Triton X-100 for 10 min and then blocked with blocking buffer for 1 hr at 37°C. The cover slips were blocked with Duolink Blocking Solution in a pre-heated humidified chamber for 30 min at 37°C. The primary antibodies to detect BRD9 (Rabbit) and FLAG (Mouse) were added to the coverslips and incubated overnight at 4°C. Then coverslips were washed with 1X Wash Buffer A and subsequently incubated with the two PLA probes (1:5 diluted in antibody diluents) for 1 hr, then the Ligation-Ligase solution for 30 min, and the Amplification-Polymerase solution for 100 min in a pre-heated humidified chamber at 37°C. Before imaging, slides were washed with 1X Wash Buffer B and mounted with a cover slip using Duolink In Situ Mounting Medium with DAPI. Fluorescence images were acquired using a Leica TCS SP8 confocal microscope. The signal was detected as a distinct fluorescent dot and analysed by fluorescence microscopy. Negative controls consisted of samples treated as described above but with only primary, secondary or control IgG antibodies.

### Generation of ARID1A knockout cells

For the generation of stable ARID1A KO cells, AD-293 cells were transfected with the NickaseNinja (ATUM) vector (pD1401-AD: CMV-Cas9N-2A-GFP, Cas9-ElecD) co-expressing two gRNAs targeting ARID1A. ARID1A gRNA sequences (GGGGAGCTCAGCGCGTAGGC and TGGCACTCCGGGCTCCGGCG) were designed using the ATUM gRNA Design Tool. 48 hrs post-transduction, positive GFP cells were sorted by FACS (BD FACSAriaTM Fusion) and plated as single cells into 96-well plates. After 15 days, clones were collected, expanded and validated by western blotting.

### Generation of inducible DBTRG-05MG cell cultures and multiciliogenesi**s**

GEMC1 and MCIDAS were amplified from FLAG-hBirA*-GEMC1/MCIDAS using forward primers and reverse primers containing *Spe*I-*Xho*I restriction sites (GEMC1 F– AAAAACTAGTatggactacaaagacgatgac, R- TTTTCTCGAGCTAAGACTGCTTAGGGACCCA), (MCIDAS, F– AAAAACTAGTatggactacaaagacgatg, R- TTTTCTCGAGTCAACTGGGGACCCAGCGGAAC) using KOD Hot Start DNA Polymerase (Millipore) and cycling conditions recommended from the manufacturer (polymerase activation at 95 °C for 2 min, denaturation at 95 °C for 20 s, annealing at 55 °C for 10 s and extension at 70 °C for 50 s, repeated for 40 cycles). PCR products were purified using the PureLink Quick Gel Extraction Kit (Invitrogen) and ligated into the pENTR vector for final cloning into the pSLIK-Neo destination vector (Addgene 25735)^23^ using LR Clonase gateway reaction vector (Invitrogen). Top10 competent *E. coli* cells (Invitrogen) were transformed and colonies were selected with neoomycin. Constructs were verified by restriction digestion and sequencing (Macrogen). The pSLIK-Neo-FLAG-hBirA*-GEMC1/MCIDAS plasmids were co-transfected with lentiviral packaging plasmid vectors REV (Addgene 12253)^24^, RRE (Addgene 12251)^24^ and VSV-G (Addgene 8454)^25^ into AD-293 cells with PEI (Sigma-Aldrich). Two days after transfection, virus-containing medium was collected and filtered through a 0.45-µm low-protein-binding filtration cartridge. The virus containing media was directly used to infect DBTRG-05MG cells (obtained from DSMZ) in the presence of polybrene (8 µg/mL) for 48 hours, before 400 μg/ml Neomycin (G418) was introduced for 72 hours. Recovered cells were then tested for successful introduction of protein expression and induction via western blot. For induction experiments, expression was induced on Day 0 by addition of Doxycycline 1 μg/ml with the addition of 5 μM DAPT (Sigma). The next day media was changed to 1% FBS whilst maintaining Doxycycline and notch inhibition. On Day 3 the media was replenished with media containing Dox and DAPT with the addition of Noggin 20 ng/ml (IRB Barcelona protein production facility). Measurements were taken at Days 0, 3 and 7 from cells plated on coverslips. Immunostainings were performed as previously described, using antibodies listed in Supplementary Table S1. dBRD9 (Tocris) was applied at Day 0 in conjunction with Dox induction. PLA was performed as previously described and then slides were counter stained with Acetylated tubulin before mounting. Fluorescence images were acquired and denconvolved using a Leica TCS SP8 confocal microscope.

### MTEC ALI cultures

All animal studies were performed following protocols that are compliant with guidelines of the Institutional Animal Care and Use Committee at Washington University and the National Institutes of Health. Mouse Tracheal Epithelial Cell (MTEC) cultures were established as previously described^26,27^. Briefly, C57BL/6J mice were euthanized at 2-4 months of age, trachea were excised, opened longitudinally to expose the lumen, placed in 1.5 mg/ml pronase E in DMEM/F-12 supplemented with antibiotics and incubated at 4°C overnight. Tracheal epithelial progenitor cells were dislodged by gentle agitation and collected in DMEM/F12 containing 10% FBS. After centrifugation at 4°C for 10 min at 800 g, cells were resuspended in DMEM/F12 with 10% FBS and plated in a Primaria Tissue Culture dish (Corning) for 3-4 hrs at 37°C with 5% CO2 to adhere contaminating fibroblasts. Non-adherent cells were collected, concentrated by centrifugation, resuspended in an appropriate volume of MTEC-complete medium (described in^28,29^), and seeded at a density of 9X10^4^ cells/cm^2^ onto Transwell-Clear permeable filter supports (Corning) treated with 0.06 mg/ml rat tail Collagen type I. Air liquid interface (ALI) was established after cells reached confluence by feeding cells with MTEC serum-free medium^28,29^ only in the lower chamber. Cells were treated with either 5, 10 or 20 μM I-BRD9 in serum-free medium, which was exchanged every 2 days. Cells were cultured at 37°C with 5% CO2, and media containing I-BRD9 replaced every 2 days for up to 12 days. Samples were harvested at ALI days 4, 8, and 12, fixed in either 100% ice-cold methanol or 4% paraformaldehyde in PBS at room temperature for 10 min.

### Statistics

Statistical significance was evaluated using two-sided paired or unpaired *t*-tests, one way Anova or a generalized linear model as was appropriate for the individual experiment and indicated in the figure legends. In the figures with bar graphs, bar indicates the mean ± S.D.

## Results

### GEMC1 and MCIDAS activate common and specific target genes

Transient overexpression of either GEMC1 or MCIDAS is sufficient to activate key genes implicated in MCC differentiation, including *TP73, MYB, CCNO* and *FOXJ1*, in a manner dependent on the coiled-coil (CC) and C-terminal TIRT domain of either protein^5,6,12,13,16^. We took advantage of this facile system to try and understand their relative influence on gene expression. Microarray analysis of gene expression changes in AD-293 cells expressing either GEMC1 or MCIDAS following transient transfection, revealed the activation of a similar core set of genes implicated in MCC differentiation (Figure 1A, B and Supplementary Table S2). This included several transcription factors important for MCC differentiation, including *FOXJ1, TP73, MYB* and *FOXN4*, as well as additional genes implicated in the MCC program, such as *CCNO* and *CDC20B*. GEMC1 and MCIDAS expression led to the differential expression (p<0.05, fc ≥ 0.25 or ≤-0.25) of a similar number of genes, and around 50% of those induced by GEMC1 were common to MCIDAS (Figure 1C). Functional enrichment analysis on gene ontologies (GO) revealed that GEMC1 and MCIDAS activated genes in overlapping GO categories, including Ribosome biogenesis, Developmental maturation, Transcription factor binding and Cilium, with the latter only significantly enriched by MCIDAS (Figure 1D and Supplementary Table S3). Despite the similarity in their transcriptional profiles, both proteins also activated some targets preferentially (Figure 1C). Several genes were highly enriched with *MCIDAS* in comparison to *GEMC1*, including *CCDC96* and *HSPA1L,* that have both been previously implicated in centrosome or cilium biology (Figure 1B)^30–32^.

### Identification of the proximal interactomes of GEMC1 and MCIDAS

As neither GEMC1 nor MCIDAS contain recognizable DNA binding domains, we wanted to better understand how they regulate transcription and achieve distinct target specificity. To address this, we performed BioID-mass spectrometry (BioID-MS) to identify proximal interactors for each protein. Both genes were N-terminally tagged with a FLAG-tagged BirA* enzyme and expressed in AD-293 cells supplemented with biotin for 20 hours (Figure 2A and Supplementary Figure S1). Biotinylated proteins were affinity purified, trypsinized and identified by MS. The proximal interactomes of GEMC1 and MCIDAS were highly similar and the proteins identified were almost exclusively involved in transcription (Figure 2B, C and Supplementary Table S4). Validating the approach, known interactors of each protein were identified (Figure 2C, D). This included Geminin, that interacts with the coiled coil (CC) domain of both proteins and its primary interactor CDT1, as well as E2F4/5-DP1 that interacts with the C-terminal TIRT domain of GEMC1 or MCIDAS^5,8,9,12^.

Consistent with previous work, MCIDAS showed increased labeling of E2F4/5-DP1 (TFDP1) compared to GEMC1 (Figure 2D)^13^, and E2F3, DP-2 (TFDP2) and the E2F regulators RB (RB1) and p107 (RBL1) were also identified. Multiple components of the DREAM/MuvB complex, including MYB-B (MYBL2), LIN9, LIN52, and LIN54, that functions with E2F to regulate cell cycle related genes, were identified with both baits (Figure 2D). TRRAP, that was previously shown to be required for multiciliation, as well as many components of its associated chromatin regulatory complexes SAGA/STAGA, ATAC and NuA4, were identified (Figure 2D and Supplementary Figure S1)^33–35^. In addition, many other transcriptional regulatory complexes and factors, including the p300 and CBP acetyltransferases, SWI/SNF, Mediator, LSD-CoREST, Polycomb (PRC1, PR-DUB), and COMPASS-like complexes were identified to different extents with both baits (Figure 2D and Supplementary Figure S1). Similarly, TP73, that is transcriptionally activated by both GEMC1 and MCIDAS and shown to be a direct interactor of GEMC1, as well as the YAP1 and TEAD1 transcription factors that interact physically and functionally with TP73, were identified with both baits (Figure 2D)^6,33,36,37^.

In contrast, the Aryl hydrocarbon receptor (AHR), that is required for *CCNO* expression and MCC differentiation at some developmental stages, and its associated proteins ARNT and AIP, as well as ELMSAN1/MIDEAS and DNTTIP1, defining components of the MiDAC complex, were identified as proximal interactors highly enriched with GEMC1 compared to MCIDAS (Figure 2D and Supplementary Figure S1)^38,39^. CDK2 and Cyclin A (CCNA2) were also highly enriched in GEMC1 transfected samples and have been shown to associate with the MiDAC complex, as well as regulate E2F4/5 and RB family members (Figure 2D)^40,41^.

While many other proteins involved in coactivator and corepressor complexes, as well as basal transcription factors, splicing regulators and RNA binding proteins, were identified (Supplementary Table S4), both samples were notably enriched for components of the SWI/SNF nucleosome remodelers and the Mediator complex, that is involved in the coordination of promoter and enhancer elements and recruitment of RNA polymerase 2 (RNAP2) to sites of transcription (Figure 2D)^42,43^. Although there were differences in specific subunits, the overall abundance and identity of Mediator subunits identified was highly similar between GEMC1 and MCIDAS (Figure 2D).

While potentially hundreds of distinct SWI/SNF subcomplexes can be formed, 3 major subcomplexes, defined by distinct DNA and chromatin binding components, have been identified: BAF, PBAF and ncBAF/GBAF, that each contain overlapping and unique subunits. In addition, specificity of SWI/SNF complexes can be dictated by the specific ATPase subunit used, either SMARCA2 (BRM) or SMARCA4 (BRG1). While GEMC1 and MCIDAS identified similar core subunits, we observed differential labeling of subunit-specific complexes and ATPase domains. In the case of GEMC1, we found primarily BAF, defined by the ARID1A/B subunits, and to a lesser extent PBAF subunits and near exclusive labeling of BRG1, a known interactor of Geminin (Figure 2D)^44^. In contrast, MCIDAS labeled the ncBAF components BICRA (also referred to as GLSTCR1) and BRD9, that were not enriched with GEMC1, as well as both BRG1 and BRM ATPase subunits (Figure 2D). These data indicated that although GEMC1 and MCIDAS both labeled Mediator to a similar extent, GEMC1 and MCIDAS preferentially associated with distinct SWI/SNF subcomplexes; namely the ARID1A containing BAF complex in the case of GEMC1 and the BRD9 containing ncBAF in the case of MCIDAS. These results established that the proximal interactomes of GEMC1 and MCIDAS are highly similar and comprised primarily of proteins involved in transcription, including many factors required for multiciliation.

### The C-terminal domains impart distinct SWI/SNF interactions

To further validate some of these results, we performed small scale BioID affinity purifications (BioID-AP) followed by western blotting in transfected AD-293 cells. We observed that expression of either GEMC1 or MCIDAS fused to BirA* led to similar, robust levels of YAP1 biotinylation that was not observed in controls (Supplementary Figure S2A). Similarly, the SMARCE1 (BAF57) component of the SWI/SNF remodeler, that plays important roles in transcriptional regulation, was labeled to a similar extent by both GEMC1 and MCIDAS, but not BirA* alone, consistent with the proteomic data (Supplementary Figure S2A).

SWI/SNF nucleosome remodeling complexes occupy thousands of genomic loci and play important roles in transcriptional regulation during development, including the regulation of E2F target genes^45^. As the enriched interactions between GEMC1 and BAF and MCIDAS and ncBAF emerged as one of the major differences, we sought to validate them and determine if they influenced the transcriptional activity of either protein. The most abundant proximal interaction with a specific SWI/SNF component we identified was between GEMC1 and ARID1A. We further examined this using BioID-AP followed by western blotting. We observed ARID1A in Strep-purified lysates from both GEMC1 and MCIDAS proteins but enriched in GEMC1 samples, consistent with the BioID-MS data (Figure 3A). In contrast, we observed BRD9, a defining component of the ncBAF complex, highly enriched in MCIDAS samples, while the levels of core SWI/SNF components SMARCE1 (BAF57) and SMARCC1 (BAF155) were similar in each case and not identified in the BirA* controls (Figure 3A).

To try and understand what region of the protein conferred the BRD9 proximal interaction, we swapped the C-termini of GEMC1 and MCIDAS directly after the CC domain to generate the hybrid proteins GEM-IDAS and MC-C1 (Figure 3B). In previous work, the C-terminal TIRT domain of MCIDAS was shown to have a higher affinity for E2F4/5-DP1 than GEMC1 when compared directly^13^, consistent with our BioID results (Figure 2D). To determine if swapping the larger C-terminal domains also influenced this higher affinity interaction, we performed immunoprecipitations for GEMC1, MCIDAS, GEM-IDAS and MC-C1 using an HA-tag. As predicted, GEM-IDAS, containing the C-terminus of MCIDAS, immunoprecipitated higher levels of both E2F4-DP1 and E2F5-DP1 than GEMC1 or MC-C1 containing the C-terminus of GEMC1 (Figure 3D and Supplementary Figure S2B).

We next examined the proximal interactions between the hybrid proteins and specific components of the BAF or ncBAF complex using BioID-AP westerns. GEM-IDAS again showed a similar profile as MCIDAS, interacting proximally with BRD9, and to a lesser extent with ARID1A. In contrast, MC-C1 did not exhibit detectable labeling of either specific component and showed reduced labeling of the core SMARCE1 component, despite showing similar levels of overall expression (Figure 3D).

Previous work reported antagonistic interactions between Geminin and BAF that required an unstructured region C-terminal to the CC of Geminin^44^. While this region is not clearly conserved in GEMC1 or MCIDAS, we sought to determine if the specificity for BRD9 was influenced by residues in the disordered region between the CC and TIRT domains. To address this, we generated 2 additional hybrid constructs; GEM-IDAS-C1 that contained the protein sequence of MCIDAS from between the CC and TIRT, inserted into GEMC1, and GEM-IDA-C1, that included the same sequences with a deletion of a possibly structured MCIDAS domain, predicted by PONDR (www.pondr.com), within the region derived from MCIDAS (Figure 3B and Figure S3)^46^. Both of these constructs were tested in BioID westerns, and despite an intact GEMC1 N-terminus, CC and TIRT domain, no interactions with either ARID1A or BRD9 were detected when the intervening sequences from MCIDAS were present (Supplementary Figure S2C). This indicated that in the context of the CC and TIRT domains of GEMC1, the maintenance of this region is critical for efficient BAF/ARID1A complex interactions. However, the insertion of the corresponding MCIDAS region was not sufficient to alter the specificity towards ncBAF binding. Finally, we created another hybrid protein, MC-GEM-S, that contained the N-terminus of MCIDAS until the end of the CC, as well as the MCIDAS TIRT domain, with the intervening sequence from GEMC1 inserted between them. This construct efficiently labeled BRD9, indicating that the distinct MCIDAS TIRT domain was able to facilitate ncBAF interactions and that the additional MCIDAS sequence between the CC and TIRT, including the predicted structured region, was not required for the proximal interaction (Figure 3E).

To validate these results further in cells, we used the Proximity ligation assay (PLA) to examine the subcellular localization of GEMC1, MCIDAS or the hybrid proteins, all tagged with FLAG, relative to BRD9 (Figure 3F and 3G). We observed that all of the FLAG-tagged proteins localized to the nucleus, in some cells forming discrete foci or localizing diffusely throughout the nucleus (Figure 3G, lower panels). In PLA experiments, we observed signal generated by the proximity of the FLAG and BRD9 antibodies only in cells transfected with constructs containing the MCIDAS TIRT domain (MCIDAS, GEM-IDAS and MC-GEM-S) consistent with the specific interaction of MCIDAS observed in BioID-MS (Figure 2D), as well as BioID-AP results (Figure 3A, 3D and 3E). Together these results demonstrated that the TIRT domain of MCIDAS imparts its specific interactions with BRD9 containing ncBAF SWI/SNF complexes and indicate that both the N and C-terminal portions of GEMC1 are required for its efficient interactions with ARID1A containing BAF complexes.

### Distinct SWI/SNF complexes influence transcriptional activation

Given the clear proximal interactions with SWI/SNF and their established role in E2F-mediated transcription, we sought to determine if they were required for GEMC1 or MCIDAS-mediated transcriptional activation. We generated AD-293 cells lacking ARID1A using CRISPR/CAS9 (Figure 4A) and analyzed the ability of GEMC1 or MCIDAS to activate several common or enriched gene targets. In *ARID1A-KO* cells, neither protein could activate *FOXJ1* or *CCNO* following transfections (Figure 4B). In contrast, the activation of several genes preferentially activated by MCIDAS (Figure 4C), *CCDC96* and *HSPA1L,* were largely unaffected by ARID1A status (Figure 4C). Conversely, the activation of both of these genes was reduced following treatment with a small molecule inhibitor of BRD9 (I-BRD9) (Figure 4D)^47^. Treatment with I-BRD9 also strongly reduced the activation of *FOXJ1* by MCIDAS but had a minor effect on its activation by GEMC1 (Figure 4E).

**Figure 4.**
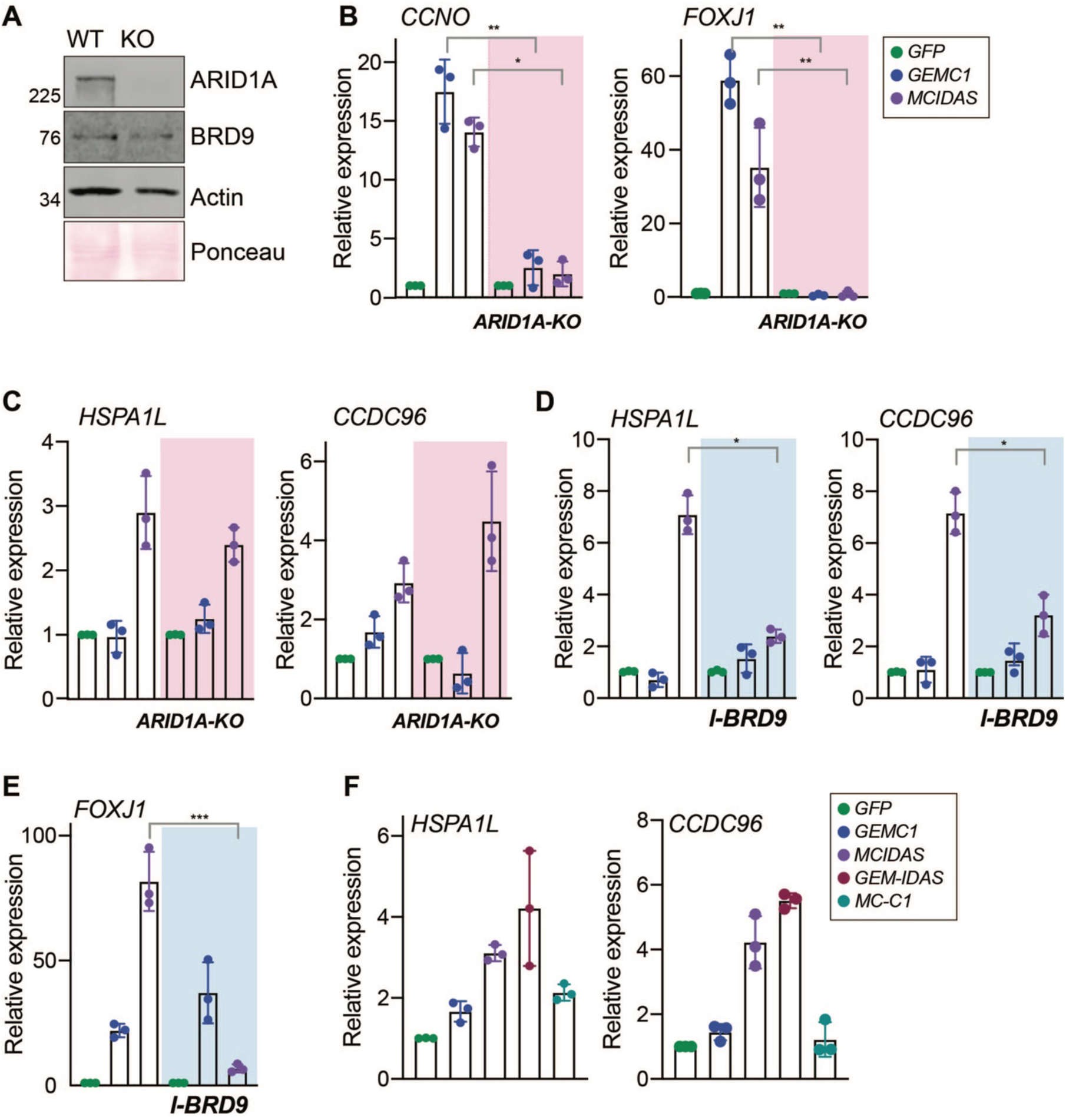
Transcriptional dependence on SWI/SNF subcomplexes. **A.** Validation of AD-293 ARID1A knock-out (KO) cells. Western blots for ARID1A and BRD9 are shown. Actin and Ponceau staining serve as loading and transfer controls. **B.** Quantitative real-time PCR (qRT-PCR) analysis of *CCNO* and *FOXJ1* expression in AD-293 cells following transfection with the indicated genes. WT vs ARID1A-KO p-values were 0.0027** and 0.0083** for *GEMC1* and 0.0142* and 0.0024** for *MCIDAS*. Transfected genes are color coded in the key to the right. **C.** qRT-PCR analysis of *HSPAL1* and *CCDC96* in parental or ARID1A-KO AD-293 cells. Statistical analysis as in B revealed no significant differences. **D.** qRT-PCR analysis of *HSPAL1* and *CCDC96* in parental or ARID1A-KO AD-293 cells treated with 5 μM of BRD9i. WT vs BRD9i p-values were 0.0207* (*HSPA1L*), 0.0166*(*CCDC96*) and 0.0176*(*FOXJ1*) for *MCIDAS*. Color coding as shown in panel B. **E.** qRT-PCR analysis of FOXJ1 expression in AD-293 cells following transfection with the indicated genes, p=0.0003*** for *MCIDAS*. Color coding as shown in panel B. **F.** qRT-PCR analysis of *HSPAL1* and *CCDC96* expression in AD-293 cells following transfection with the indicated genes. Transfected genes are color coded in the key to the right. For B-F, results from 3 independent experiments are shown with the bars indicating the mean and the standard deviation shown. For statistical analysis a paired (B, C) or unpaired (D, E, F) t-test was used, p≤0.001=***, p≤0.01=** and p≤0.05=*.

As replacing the C-terminus of GEMC1 with that of MCIDAS resulted in a proximal interaction with BRD9, we analyzed the ability of the hybrid proteins to activate *HSPA1L* and *CCDC96*, that were ARID1A independent, but sensitive to I-BRD9. The GEM-IDAS fusion protein, that showed BRD9 interactions (Figure 3D), activated both genes to a similar extent as MCIDAS, while the MC-C1 fusion protein showed only minor activation, similar to that of GEMC1 (Figure 4F). Together, these results demonstrated that ARID1A and BRD9 influence the transcriptional output of GEMC1 and MCIDAS. In addition, they predicted that ARID1A (BAF) or BRD9 (ncBAF) loss may differentially affect subsets of GEMC1 or MCIDAS targets and impair multiciliogenesis.

### BRD9 depletion inhibits multiciliogenesis in an inducible human cell model

To determine if BRD9 inhibition influenced multiciliation, we established a new inducible system to generate MCCs in human cells. We transduced cultures of the human glioblastoma cell line DBTRG-05MG with Doxycycline inducible expression vectors for FLAG tagged GEMC1 or MCIDAS. Following 3 days of serum starvation, Dox treatment was used to induce the transgene (Day 0) and cultures were treated with the NOTCH inhibitor, DAPT (Figure 5A). At Day 0, all cells had one or no cilia, visualized by acetylated tubulin staining (Figure 5B and 5C). Cells were treated with the WNT inhibitor noggin at Day 3, at which time there was a significant amplification of centrioles marked by centrin staining in FLAG positive cells, as well as corresponding acetylated tubulin, indicating basal body formation (Figure 5A, 5B and 5C). FLAG expression showed that GEMC1 and MCIDAS both localized to the nucleus and this was unaffected by treatment with the BRD9 PROTAC dBRD9 (Figure 5B, left panels). Treatment with dBRD9 inhibited the amplification of centrioles seen at Day 3 in the MCIDAS inducible system (Figure 5B and 5C). At Day, 7 multiple cilia were evident with more than 10 cilia in almost 50% of the FLAG positive cells expressing MCIDAS or GEMC1 (Figure 5B and 5C). Consistent with the impaired centriole amplification, treatment with dBRD9 almost completely prevented the production of multiple cilia in MCIDAS expressing cells (Figure 5B, bottom row and 5C).

**Figure 5:**
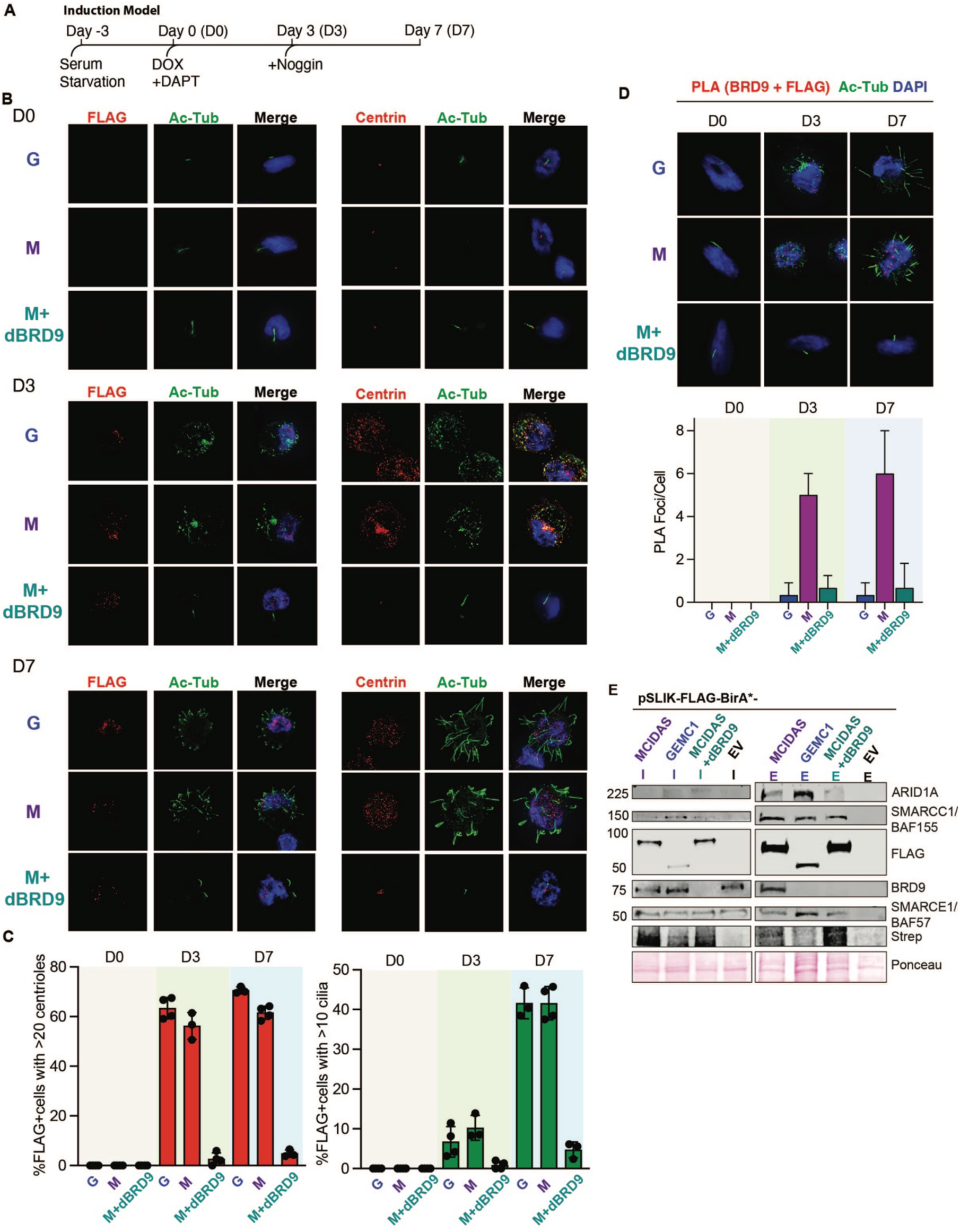
BRD9 depletion inhibits multiciliogenesis in a cell line model. **A.** Schematic demonstrating the time course of the inducible system used to generate MCCs in human DBTRG-05MG cells. For full details, see methods section. **B.** Immunofluorescence using the indicated markers in cells expressing either G=pSLIK-FLAG-BirA*-GEMC1, M=pSLIK-FLAG-BirA*-MCIDAS or M+DBRD9=pSLIK-FLAG-BirA*-MCIDAS+dBRD9 (20uM). Images are taken at three different time points Day 0, Day 3 and Day 7. **C.** Quantification of Immunofluorescence shown in B. Centriole and cilia counts were generated in FLAG positive cells, using Centrin and acetylated tubulin (Ac-Tub) as markers. N=3 or 4 independent experiments with a total of 150-200 cells scored. Averages of each experiment are shown (ovals) with mean (vertical bars) and standard deviation. **D**. Proximity ligation assays (PLA) counter stained with Ac-Tub and DAPI, corresponding to the conditions in **E**. Quantification of PLA foci are shown in bottom panel. N=100 cells per condition. **F**. BioID-AP followed by western blotting at Day 3. I indicates Input and E indicates Eluate. I and E westerns run separately, but from same BioID-AP sample.

Using the same system, we applied PLA to visualize the proximal interaction between BRD9 and either GEMC1 or MCIDAS (using FLAG) and the effects of dBRD9 on this interaction (Figure 5D). At Day 3, PLA with FLAG (MCIDAS) and BRD9 antibodies resulted in nuclear foci representing their proximal interaction (Figure 5D). This strong increase in foci was not observed in cells expressing GEMC1, or in MCIDAS expressing cells treated with dBRD9 (Figure 5D). At Day 7, when multiple cilia were observed, similar proximal interactions between MCIDAS and BRD9 were detected with little signal in GEMC1 expressing or dBRD9 treated cells (Figure 5D). Finally, we performed a BioID-AP western at the Day 3 time point and found that dBRD9 efficiently degraded BRD9, thus abolishing the interaction in BioID-AP western (Figure 5E). These results further support a role for SWI/SNF complexes in multiciliogenesis and indicate that BRD9, and by extension ncBAF complexes, are crucial regulators of multiciliogenesis in this system.

### Inhibition of BRD9 inhibits multiciliogenesis

To further test whether SWI/SNF complexes were important for MCC differentiation in a more well established system, we used mTECs that can generate MCCs and other differentiated cell types characteristic of the airways when exposed to ALI^27^. ALI cultures were treated with I-BRD9 during differentiation at 3 doses and examined for their ability to generate MCCs. Treatment with I-BRD9 reduced the expression of both *Foxj1* and *Trp73* at the mRNA and protein level in a dose-dependent manner (Figure 6A-6C). In addition, there was a dose-dependent reduction in cells exhibiting centriole amplification (inferred by staining for Centrin and Deup1) (Figure 6A, D, E) and multiciliogenesis (inferred by staining with Acetylated tubulin antibodies (Ac-tubulin) (Figure 6A, F). Together these data indicated that ncBAF function is essential for multiciliogenesis in murine ALI mTEC cultures.

**Figure 6:**
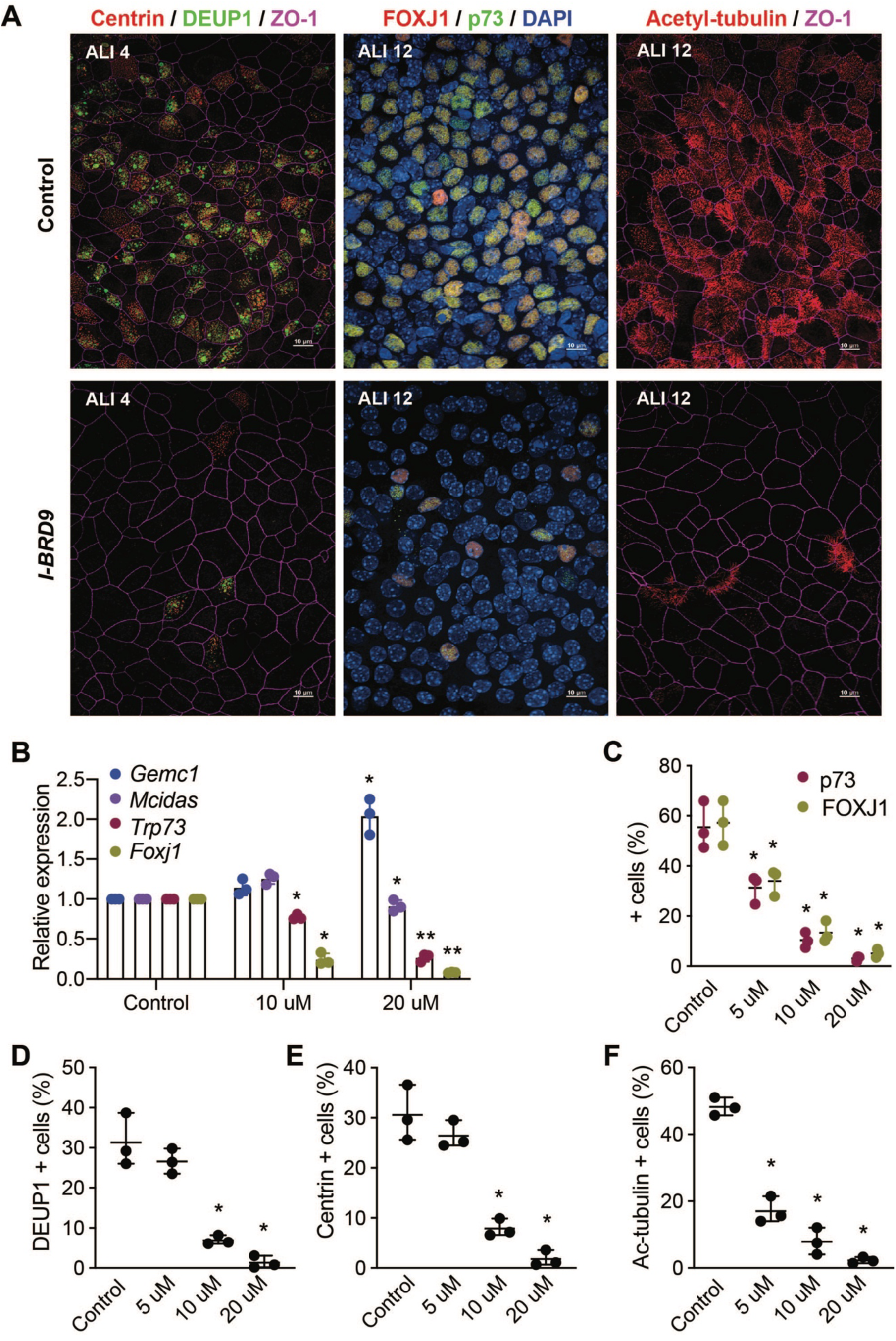
BRD9 inhibitors impair multiciliation in MTECs. **A.** Representative images of MTEC ALI cultures stained for Centrin and Deup1 to analyze centriole amplification, Foxj1 and p73 transcription factor targets of GEMC1/MCIDAS, or Acetylated tubulin (Ac-tubulin) to monitor ciliogenesis. Some images stained with ZO-1 to define cell boundaries or DAPI to define the nuclear DNA. ALI culture day and treatment with 20 uM BRD9i indicated. **B.** Quantitative real-time PCR analysis of *Gemc1, Mcidas, Trp73* and *Foxj1* in ALI cultures. Results from 3 independent experiments are shown with bar indicating mean and standard deviation. Paired t-tests were used for statistical analysis: Control vs. BRD9i: *Foxj1* p=0.0125* (10 uM) and 0.003** (20 uM), *Trp73* p=0.0092* (10 uM) and 0.0087** (20 uM), Gemc1 p=0.0082** (20 uM). **C.** Fraction of cells committed to the MCC fate (p73+ or FoxJ1+) at ALI 12. Results are from 3 independent experiments. N (total cells) = control (2128), 5 uM (1665), 10 uM (1273), 20 uM (1360). **D.** Fraction of cells undergoing centriole amplification (centrin-positive cells) or **E.** deuterosome formation (Deup1-positive cells) at ALI 4. Results are from 3 independent experiments. N (total cells) = control (1269), 5 uM (1062), 10 uM (1199), 20 uM (1170). **F.** Fraction of ciliated cells at ALI 12. Results are from 3 independent experiments. N (total cells) = control (1814), 5 uM (1295), 10 uM (1113), 20 uM (1099). Statistical analyses for C-F were done using one-way ANOVA. *denotes p<0.05.

## Discussion

MCCs have an intriguing developmental program in which post-mitotic MCC precursors support the explosive generation of numerous centrioles that mature into basal bodies for multiciliation. In addition, the direct involvement of MCC dysfunction in the etiology of a broad spectrum of human pathologies like respiratory disorders, fertility issues and hydrocephalus, has led to a significant degree of interest in the biology of this specialized ciliated cell-type. We and others have previously established that the two Geminin family members, GEMC1 and MCIDAS, are key factors that organize the transcriptional regulation of MCC differentiation^18^. Yet, several lines of evidence point to distinct functions for GEMC1 and MCIDAS in MCC differentiation that is not adequately explained by our current understanding of their molecular interactions. In this work, we compared their transcriptional targets and proximal interactions and found that, while their targets and interactomes were highly similar, they also had distinct molecular interactions and transcriptional effects. The BioID approach identified nearly every protein previously implicated in the transcriptional regulation of MCC differentiation to date, including E2F4/5-DP1, TRRAP, AHR, MYB, TP73, and CDK2, as well as many additional proteins implicated in their regulation, demonstrating the suitability of this strategy for interrogating transcriptional complexes^18^. These results also indicated that the proposed role of GEMC1 and MCIDAS in DNA replication is likely limited to their interactions with Geminin/CDT1, as nearly all of the proteins identified in BioID were related to transcriptional regulation^2,7^.

Previous work showed that the E2F4/E2F5-DP1 transcription factors are required for MCC differentiation in every tissue where MCCs are present^9,48–51^. Our data suggests that E2F3, as well as DP-2 (TFDP2), may also influence the transcriptional response, particularly with MCIDAS, that has higher affinity for E2F4 and E2F5 through its distinct C-terminal TIRT domain ^13^. Associations of GEMC1 and MCIDAS with the DREAM/MuvB complex are also potentially relevant, as this complex functions with the E2F program to control cell cycle-specific transcriptional programs^18,52^. Finally, we identified RB and p107 of the pocket protein family, that presumably must be removed from repressive E2F4/5 complexes to activate transcription of their target genes^41^. GEMC1 specifically associated with CCNA2/CDK2, that are known to phosphorylate and dissociate RB/p107 during normal cell cycle (Figure 2). An attractive hypothesis is that the delivery of CDK2 activity to RB/p107 via E2F-GEMC1-TIRT domain interactions could facilitate this activation step.

Multiciliated cells utilize canonical cell cycle regulators, including CDK2, CDK1/CCNB1, PLK1 and APC/C^CDC20^, as well as MCC-specific paralogs of cell cycle regulators, such as CCNA1, CCNO and CDC20B, to facilitate centriole amplification, although their precise roles remain largely unclear^14,17,53–57^. GEMC1 was shown to be a target of CCNA2/CDK2 kinase activity^7^ and previous work suggested that GEMC1 was negatively regulated by the poorly characterized Cyclin, CCNO (Cyclin O), potentially connecting CDK activity to its transcriptional functions^56^. We also note that several G2/M specific factors, including PLK1 and CDC20 (a component of the APC/C), proximally associated with MCIDAS but not GEMC1 (Supplementary Figure S4). MCIDAS promotes the nuclear localization of GEMININ, and both GEMININ and MCIDAS are targets for mitotic degradation^2,58^. In the case of GEMININ, and likely MCIDAS, this requires APC/C^CDC20 58^. The emerging picture suggests that GEMC1 and MCIDAS are both likely targets of cell cycle-specific regulation, providing an elegant mechanism to facilitate their stepwise actions and fine tune transcriptional output with massive centriole amplification.

Recent biochemical work established the major SWI/SNF subcomplexes and demonstrated that their DNA binding and activity is largely dictated by histone modifications^59,60^. Previously, a disordered, acidic region of GEMININ, C-terminal to the CC domain, was shown to facilitate interactions with BRG1 (SMARCA4), an ATPase that can be a component of any of the SWI/SNF subcomplexes^44^. Swapping the GEMC1 and MCIDAS C-terminal domains revealed that the sequence that confers specificity for ncBAF lies within the TIRT domain of MCIDAS (Figure 4, 5). GEMC1 showed almost exclusive interactions with BRG1 containing complexes, compared to MCIDAS that labeled both BRG1 and BRM (SMARCA2) (Figure 2). However, moving the C-terminus of GEMC1 to MCIDAS (MC-C1) eliminated interactions with complex-specific subunits, indicating additional intricacy in the SWI/SNF interactions. The predicted structures of GEMC1 and MCIDAS generated by Alphafold appear divergent in the E2F4/5-DP1 binding TIRT domain, and in the case of MCIDAS, additional structured domains are predicted, including one directly upstream of the TIRT domain. Our analysis with a large panel of hybrid proteins indicated that this domain was not required to influence TIRT-dependent interactions with BRD9 containing ncBAF complexes, although it could still potentially influence protein function and should be investigated further in future work.

ARID1A containing BAF complexes bind more predominantly to H3K4 monomethylated enhancer regions, while ncBAF binds preferentially to H3K4-trimethylated promoter regions^61,62^. As both GEMC1 and MCIDAS identified multiple SWI/SNF complexes, one possibility is that they work together at some of the same genes through distinct enhancer/promoter interactions. This is supported by the identification of common transcription factors that bind enhancer and promoter elements with both proteins in BioID experiments, as well as their similar labeling of the Mediator complex that bridges promoter and enhancer elements^63–66^. We unfortunately cannot discriminate the direct and indirect targets of GEMC1 and MCIDAS, an issue that is confounded by the fact that GEMC1 activates *MCIDAS*, influencing the level of activation of some common targets^5,10,13^. In our hands, extensive attempts to ChIP either GEMC1 or MCIDAS to identify direct targets were unsuccessful. As neither protein has an identifiable DNA binding domain, their target gene specificity is likely dictated by multivalent protein interactions rather than specific contact with DNA motifs. Our data provide a number of leads to pursue this possibility in future work, in order to better understand the mechanisms that confer transcriptional specificity.

While the majority of proteins identified by GEMC1 and MCIDAS were common, there were several proteins aside from SWI/SNF components that were strongly enriched or exclusive with one of the baits. Two of these, ELMSAN1/MIDEAS and DNTTIP1, are defining components of the mitotic deacetylase (MiDAC) complex^39^. This complex recruits histone deacetylases 1 and 2, that were not identified in the BioID experiments, to regulate cell type-specific transcriptional programs during development and exhibits its highest activity in mitotic cells^39,67,68^. The MiDAC complex has been proposed to function in transcriptional activation in neurodevelopment through the deacetylation of H4K20^68^. Notably, cBAF and ncBAF activity was shown to be differentially regulated by acetylation of H4K20, as well as other residues in the H4 tail, indicating that the interaction of GEMC1 with MiDAC could be relevant to a transition from GEMC1-cBAF to MCIDAS-ncBAF binding. While the influence of MiDAC on the transcriptional program of multiciliated cells remains unknown, it represents another interesting proximal interaction for further analysis.

We identified distinct SWI/SNF subcomplexes that influence the transcriptional outputs of GEMC1 and MCIDAS and inhibition of BRD9 impaired multiciliogenesis in 2 distinct systems. This observation may suggest direct effects on ependymal MCC differentiation in mice or human patients with SWI/SNF mutations and hydrocephaly^69,70^. Despite the limitation that our proteomic study was performed by overexpressing GEMC1 or MCIDAS in cycling AD-293 cells, our approach is strongly validated by its ability to identify most of the factors known to be involved in MCC transcription and the fact that multiple BRD9 inhibitors impair the MCC pathway. Notably absent from our proteomic results were the RFX2/3 transcription factors, that are expressed at low levels in HEK-293 (http://www.proteinatlas.org). This may explain why overexpression of GEMC1 and MCIDAS is incapable of driving efficient multiciliation in this cell type^71–76^. Consistent with this, we were able to drive multiciliogenesis in the glioblastoma cell line DBTRG-05MG, that expresses both factors (www.depmap.org). This new cell line system may help facilitate future work in defining the machinery involved in multiciliation, as it can potentially be more easily scaled up and genetically manipulated than systems relying on primary cells. In conclusion, our work provides new insights into the transcriptional machinery used by GEMC1 and MCIDAS in MCC differentiation and provides resources for further investigation into the molecular functions of these non-canonical transcriptional activators.

## Acknowledgements

We thank F. Guillemot, C. Lynch, M. Serrano and S. Brody for antibodies, F. Supek for cells and reagents, A. Holland and C. Jewett for discussions, T. Dantas for sharing unpublished data and support from the IRB Functional Genomics and Biostatistics/Bioinformatics Core Facilities.

## Author contribution statement

M.L. B.T, P.A.K, T.C, H.L, J.S, I.G.C, J.Q, G.G.G, G.P and H.Z. designed and performed experiments and generated key reagents. E.C. and B.R. processed samples, performed LC-MS/MS and analyzed MS data. M.L. B.T, P.A.K, T.C, H.L, J.S, J.Q, G.G.G, G.P, V.C, S.P, H.Z, S.R, M.M, and T.H.S analyzed experimental data. C.A. analyzed array data and performed statistical analysis of applicable data. T.H.S. and M.L. wrote the manuscript with additional editing from M.M. and S.R. V.C, S.P, B.R, H.Z, X.S, S.R, M.M. and T.H.S provided funding and supervision. T.H.S. and M.L. wrote the manuscript with additional editing from M.M. and S.R. B.T. and P.A.K. contributed equally.

## Ethics statement

X.S. is a founder and scientific advisor of Nuage Therapeutics. All other authors declare no conflicts of interest.

## Data availability statement

Microarray data has been submitted to the NCBI Gene Expression Omnibus (GEO) and has been assigned the reference number GSE161197. All other data is available from the corresponding author.

