## Supplementary material for "GEMC1 and MCIDAS interactions with SWI/SNF complexes regulate the multiciliated cell-specific transcriptional program": Supplementary material.pdf

**Supplementary Information**

| **Antigen** | **Vendor** | **Cat#** | **Species** | **Dilution** | **Application** |
|---|---|---|---|---|---|
| FLAG | Sigma | F1804 | Mouse | 1:200 | IF, PLA |
| FLAG | Sigma | F7425 | Rabbit | 1:200 | Western |
| ARID1A | Sigma | HPA005456 | Rabbit | 1:100 | Western |
| BRD9 | Bethyl | A303-781A | Rabbit | 1:100 | Western, PLA |
| BAF57 | Bethyl | A300-810A | Rabbit | 1:100 | Western |
| BAF155 | Abcam | ab72503 | Rabbit | 1:100 | Western |
| Actin | Sigma | A4700 | Mouse | 1:10,000 | Western |
| HA | Santa Cruz | sc-7392 | Mouse | 1:200 | IP |
| HA | Santa Cruz | sc-805 | Rabbit | 1:4000 | Western |
| Myc | Santa Cruz | sc-789 | Mouse | 1:4000 | Western |
| Centrin | Sigma | 04-1624 | Mouse | 1:200-1:2000 | IF |
| Acetylated tubulin | Sigma | T6793 | Mouse | 1:500-1:2000 | IF |
| Deup1 | Sigma | HPA010986 | Rabbit | 1:400 | IF |
| ZO-1 | Santa Cruz | R40.76 | Rat | 1:1000 | IF |
| p73 | Abcam | ab40658, | Rabbit | 1:200 | IF |
| FOXJ1 | Kind gift from S. Brody | - | Rabbit | 1:400 | IF |
| **Secondaries** | **Vendor** | **Cat#** | **Species** | **Dilution** | **Application** |
| anti-Mouse IgG (H+L) Alexa Fluor 680 | Thermo Fisher | A-21057 | Goat | 1:500 | IF |
| anti-rabbit IgG (H+L) Alexa Fluor 680 | Thermo Fisher | A-21076 | Goat | 1:500 | IF |
| anti-Mouse IgG2b, Alexa Fluor-488 | Thermo Fisher | A-21141 | Goat | 1:500 | IF |
| anti-Mouse IgG2a, Alexa Fluor-568 | Thermo Fisher | A-21134 | Goat | 1:500 | IF |
| anti-mouse HRP conjugate | Promega | W4028 | Goat | 1:15000 | IF |

| | | | | | |
|---|---|---|---|---|---|
| anti-rabbit HRP conjugate | Promega | W4018 | Goat | 1:15000 | IF |
| IRDye® 680LT Goat anti-Rabbit IgG Secondary Antibody | Li-COR | 925-68021 | Goat | 1:15000 | WB |
| IRDye® 680LT Goat anti-Mouse IgG Secondary Antibody | Li-COR | 926-68020 | Goat | 1:15000 | WB |
| IRDye® 800CW Goat anti-Mouse IgG Secondary Antibody | Li-COR | 926-32210 | Goat | 1:15000 | WB |
| IRDye 800CW streptavidin | Li-COR | 926-32230 | Goat | 1:10000 | WB |

**Supplementary Table S1: Antibodies used in this study.**

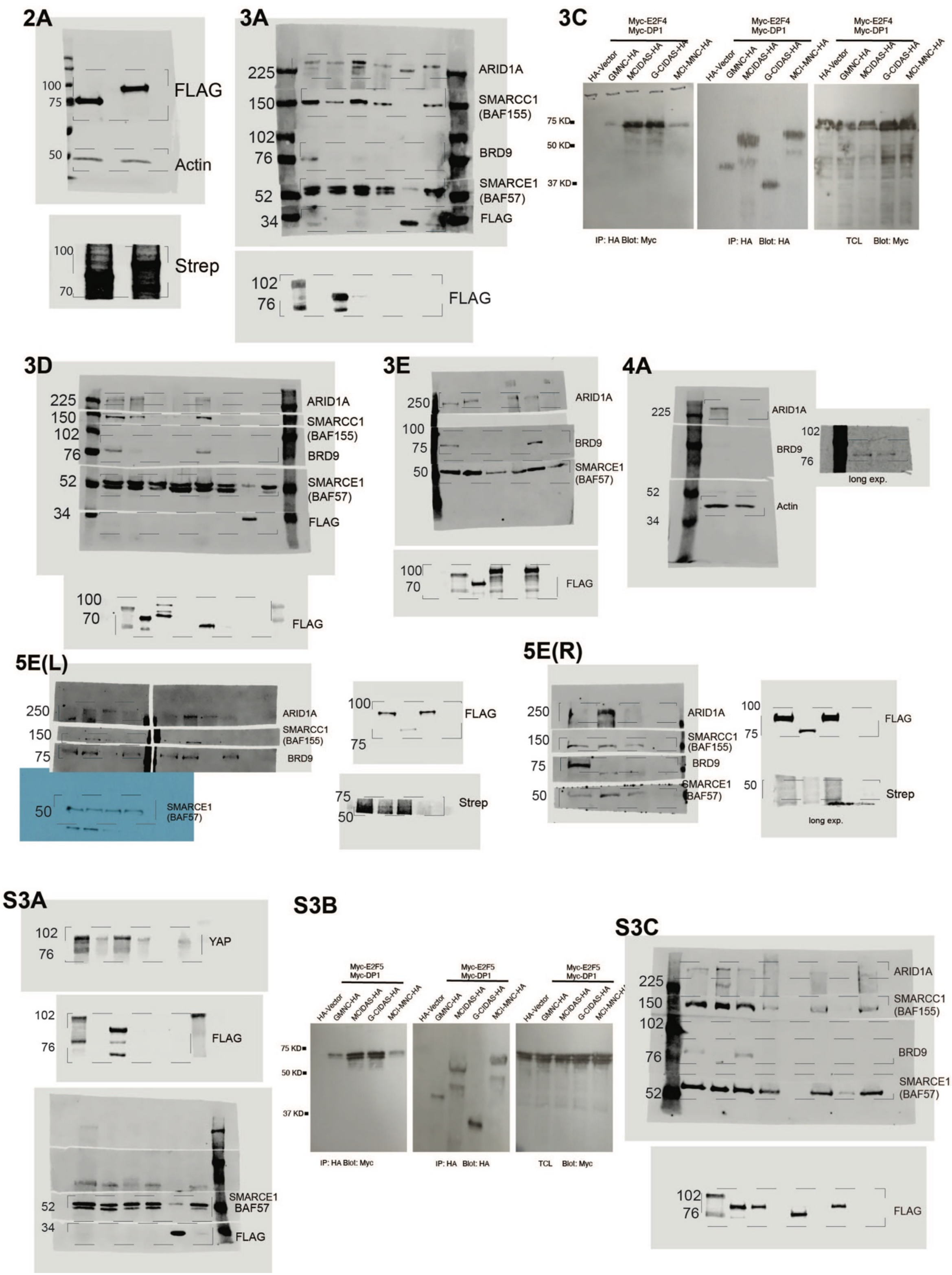


**Supplementary Figure S1: Uncropped western blots.** Figure panels, size markers and antigen are indicated. Cropping indicated by dashed lines.

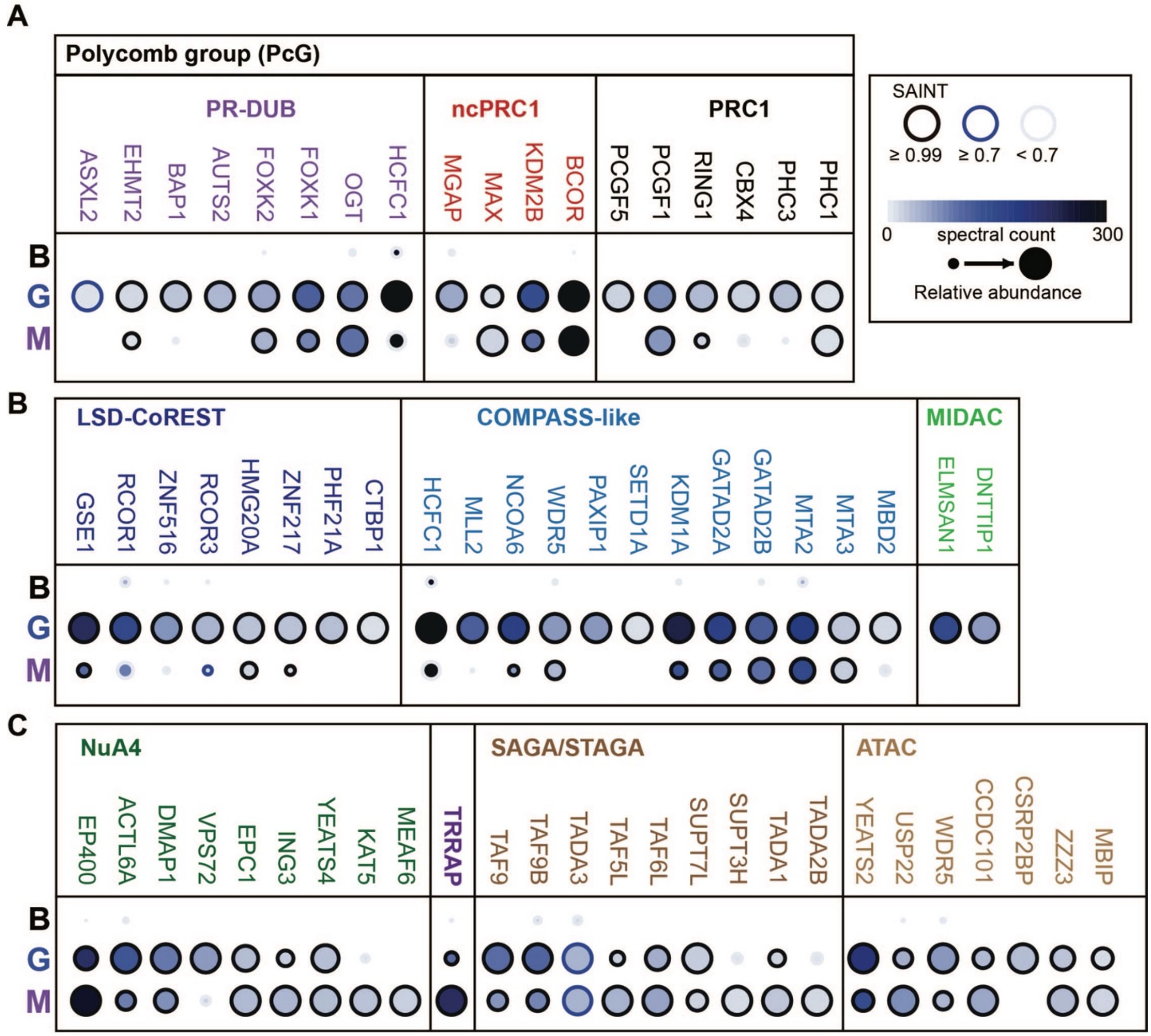


**Supplementary Figure S2: The proximal transcription factor complexes of GEMC1 and MCIDAS. A.** Dot plot depicting relative abundance and spectral counts for selected Polycomb group (PcG) proteins grouped by known complexes. Key applies to all panels. **B.** Dot plot depicting relative abundance and spectral counts for selected complexes enriched with GEMC1 compared to MCIDAS. **C.** Dot plots of transcriptional complexes associated with TRRAP. B=BirA*, G=GEMC1 and M=MCIDAS for all panels.

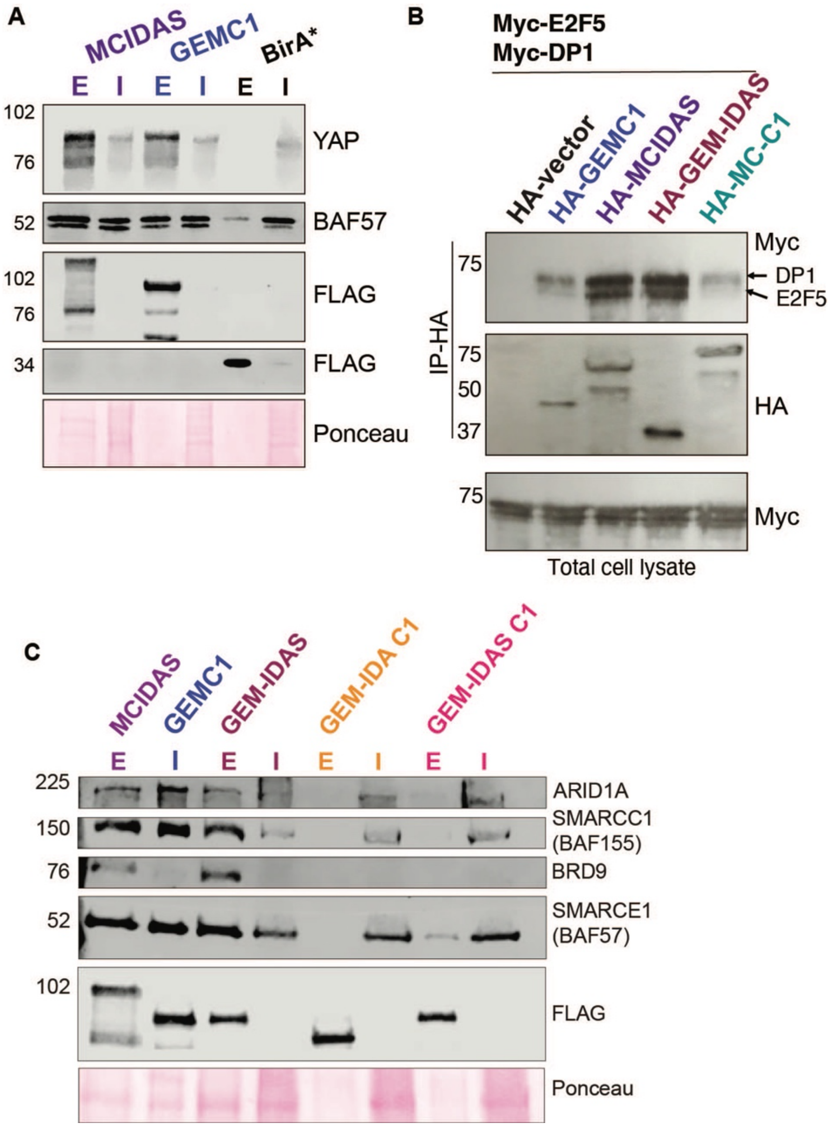


**Supplementary Figure S3: Validation of BioID-MS interactions. A.** BioID-AP western blots for YAP1, BAF57 and baits (FLAG). Input (I) and streptavidin Eluate (E) are indicated for each gene and ponceau shows similar input loading and transfer efficiencies. **B.** Co-immunoprecipitation experiments demonstrate enhanced E2F5-DP1 interactions with the MCIDAS C-terminus. HEK293T cells were transfected with HA-tagged vector, GEMC1, MCIDAS, GEM-IDAS and MC-C1, Myc-tagged DP1 and Myc-E2F5. Lysates were immunoprecipitated with anti-HA antibodies and westerns carried out for Myc and HA following transfer to PVDF. Total lysates are shown blotted for Myc. Data presented is representative of 2 biological replicates **C.** BioID-AP westerns blots for ARID1A, BRD9 and core SWI/SNF components following expression of indicated hybrid proteins (see Figure 3B for schematic). Ponceau staining shown for loading and transfer control.

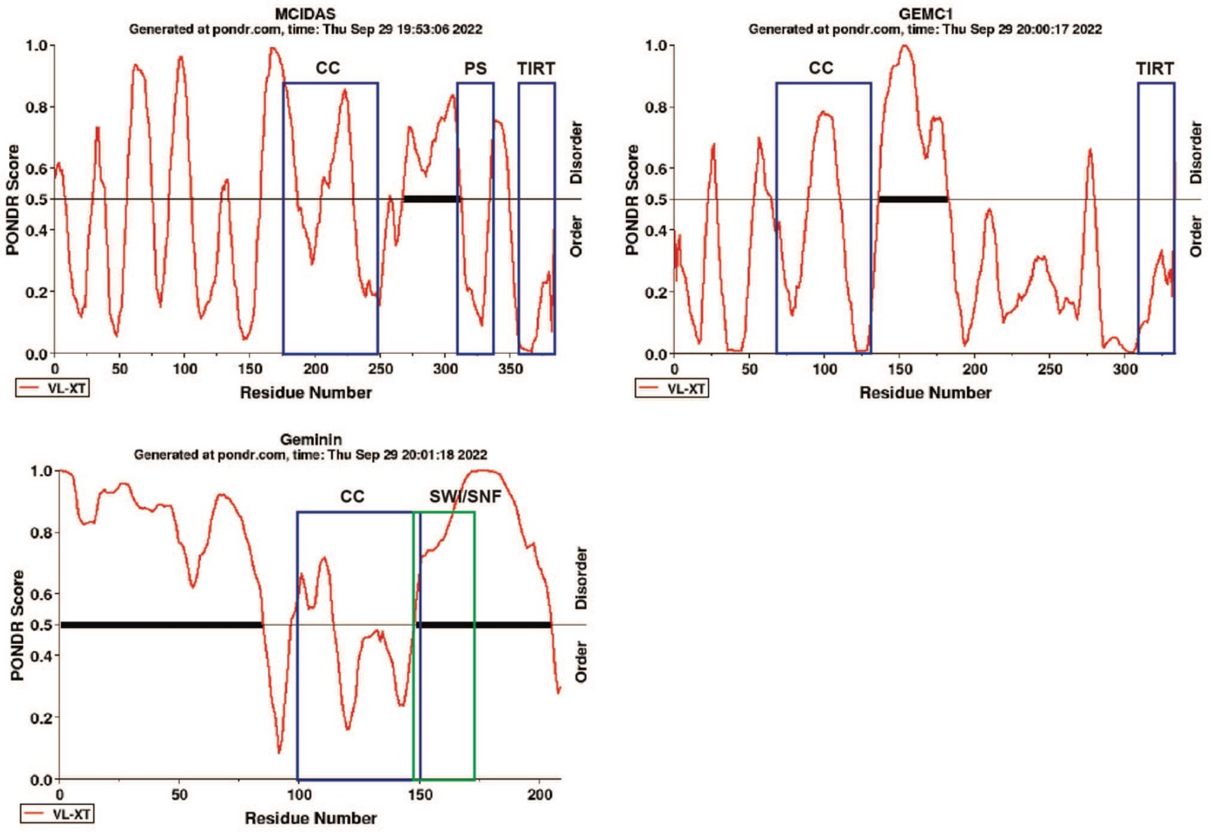


**Supplementary Figure S4: Predicted regions of disorder in Geminin, GEMC1 and MCIDAS.** Disorder of the indicated proteins predicted by PONDR. The coiled coil (CC), TIRT domains, possibly structured domain of MCIDAS (PS; see Figure 3) and SWI/SNF interacting region of Geminin are indicated.

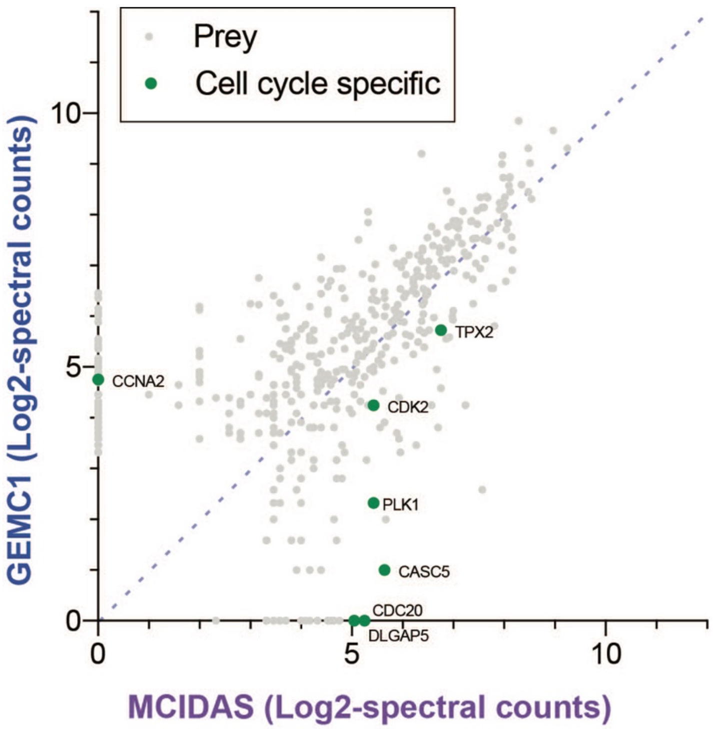


**Supplementary Figure S5. Cell cycle specific proteins identified in BioID-MS.** Scatterplot of data shown in Figure 2C highlighting proteins associated with specific cell cycle phases.
